# Deadly and venomous *Lonomia* caterpillars are more than the two usual suspects

**DOI:** 10.1101/2022.09.06.506776

**Authors:** Camila González, Liliana Ballesteros-Mejia, Juana Díaz-Díaz, Diana M. Toro-Vargas, Angela R Amarillo-Suarez, Delphine Gey, Cielo León, Eduardo Tovar, Mónica Arias, Nazario Rivera, Luz Stella Buitrago, Roberto H Pinto-Moraes, Ida S. Sano Martins, Thibaud Decaëns, Mailyn A González, Ian J Kitching, Rodolphe Rougerie

## Abstract

Caterpillars of the Neotropical genus *Lonomia* (Lepidoptera: Saturniidae) are responsible for some fatal envenomation of humans in South America inducing hemostatic disturbances in patients upon skin contact with the caterpillars’ spines. Currently, only two species have been reported to cause hemorrhagic syndromes in humans: *Lonomia achelous* and *Lonomia obliqua*. However, species identifications have remained largely unchallenged despite improved knowledge of venom diversity and growing evidence that the taxonomy used over past decades misrepresents and underestimates species diversity. Here, we revisit the taxonomy and distribution of *Lonomia* using the most extensive dataset assembled to date, combining DNA barcodes, morphological comparisons, and geographical information. Our integrative approach leads to the recognition of 60 species, of which seven are known or strongly suspected to cause severe envenomation in humans. From a newly compiled synthesis of epidemiological data, we also examine the consequences of our results for understanding *Lonomia* envenomation risks and call for further investigations of other species’ venom activities. This is required and necessary to improve alertness in areas at risk, and to define adequate treatment strategies for envenomed patients, including performing species identification and assessing the efficacy of anti-*Lonomia* serums against a broader diversity of species.

## Introduction

Caterpillars of *Lonomia* (Walker, 1855), a genus of Neotropical moths in the family Saturniidae, are among the very few lepidopterans to pose a severe threat to human health. Their venom, inoculated through spines that cover the caterpillar’s body [1], has a rather singular effect. It exhibits both coagulant and anticoagulant as well as fibrinolytic activities that can cause severe hemorrhagic syndromes, leading to diffuse hemorrhages, renal failure, brain damage and, in severe cases, death [2], [3], [4]. A *Lonomia* antivenom (LAV) is produced in Brazil against the venom of *Lonomia obliqua* (Walker, 1855) which is so far the only specific treatment available. It is used successfully across South America in cases of envenoming by *Lonomia* [5],[6],[7].

The first records of envenoming by *Lonomia* caterpillars in South America came from Venezuela in 1967, and were then attributed to *Lonomia achelous* (Cramer, 1777) [8]. Since then, several accidents have been reported in that country [9],[10], [11]. In Brazil, the Ministry of Health notified 1,930 cases between 2001 and 2006 [12]. In Colombia, the first fatal accident was reported in 2000, and the species involved was again identified as *L. achelous* [13]. Although the epidemiology of *Lonomia* envenomation in Colombia remains insufficiently documented, accidents reported to the National Institute of Health in Colombia suggest that about 10 cases occur each year. Accidents have also been reported in Peru in 2006, 2010 and 2016 [14], [15] (species unidentified), in Argentina [16],[7],[17] (all cases from Misiones province and attributed to *L. obliqua*) and in French Guiana [18],[19] (attributed to *L. achelous*).

*Lonomia* caterpillars are gregarious, forming groups of more than fifty individuals with cryptic coloration that remain inactive on tree trunks during the day (Fig. 1) and become active at night to feed on leaves of the host tree [20]. Their spines have unicellular venom glands without pores for venom liberation [21],[1]; when a spine breaks after touching or penetrating a rigid surface, such as the human skin, the venom is released [20],[21]. Incidents tend to occur when people are accidentally exposed to caterpillar colonies at rest on the tree trunks during the day [22].

**Figure 1.**
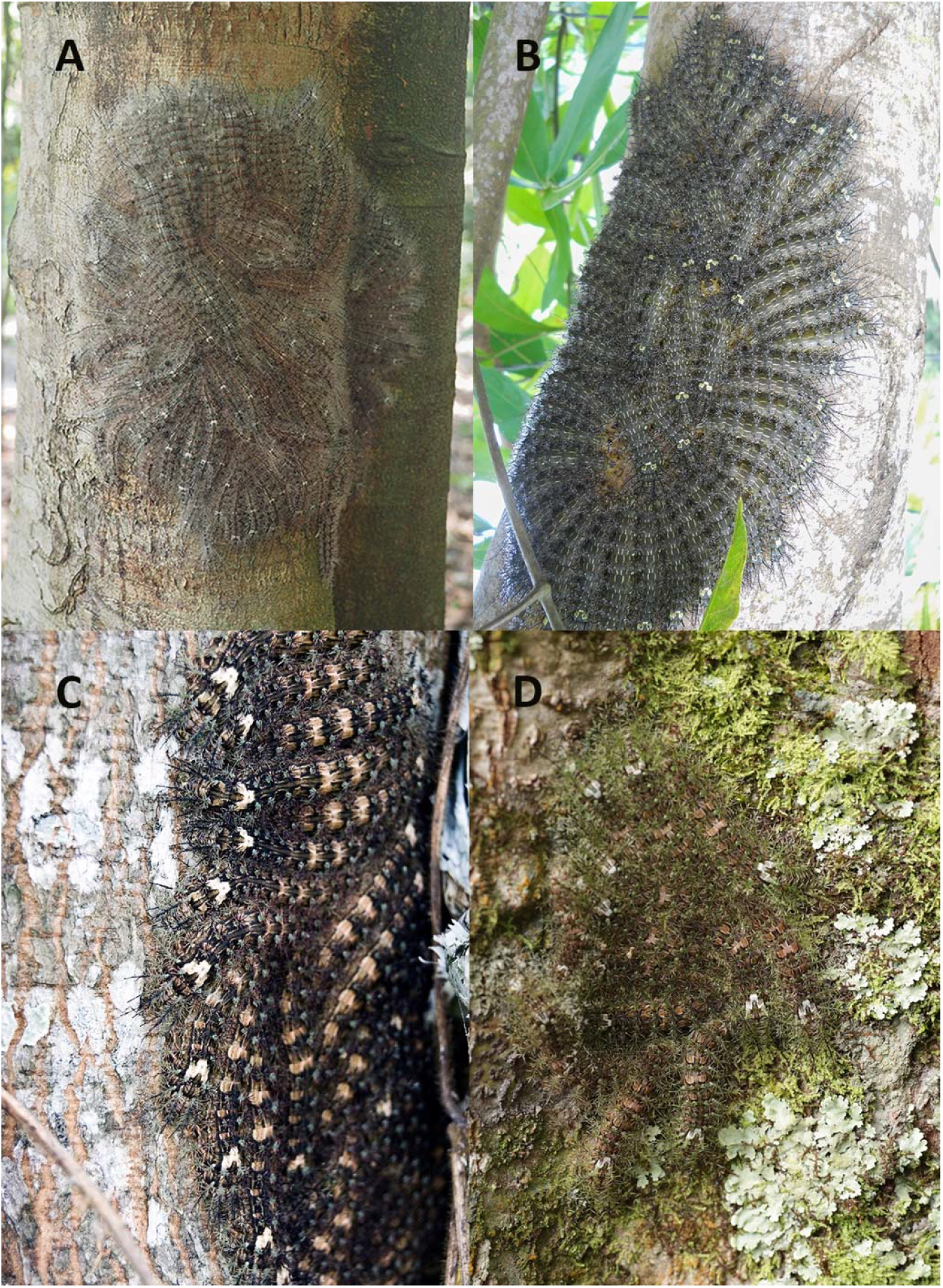
*Lonomia* caterpillar colonies. A. *Lonomia descimoni*, Leticia, Amazonas, Colombia B. *Lonomia orientoandensis*, Tauramena, Casanare, Colombia. C. *Lonomia casanarensis*, Tauramena, Casanare, Colombia. D. *Lonomia orientocordillera*, Cubarral, Meta, Colombia.

So far, when identified, only two species have been formally linked to envenomation accidents: *L. achelous* and *L. obliqua* [20]. However, [6] recently revealed the ability of two other species, namely *Lonomia orientoandensis* Brechlin and Meister, 2011 and *Lonomia casanarensis* Brechlin, 2017 to induce similar hemorrhagic syndrome in rats. This emphasizes the knowledge gap on species diversity within the genus *Lonomia* in the past 20 years. The most recent taxonomic revision [23] recognized 11 species and one subspecies of *Lonomia*, but a more recent census listed 52 valid species and 5 valid subspecies [24]. This contrast reflects the recent high rate of species discovery and description in Neotropical Saturniidae (see [24]), largely resulting from the integration of DNA barcoding [25] into taxonomic studies of these moths [26],[27]. Although all records of envenomation accidents so far have been reported to involve either *L. achelous* or *L. obliqua*, the “inflation” of the number of species in the genus and the finding that other species are also likely to be highly venomous [6] suggests that these two are unlikely to be the only species involved in cases of human envenomation and that species identification of past records might also be inaccurate in the context of the current classification. Furthermore, analyses of venom composition and physiological properties have revealed important differences between *L. achelous*, whose venom is mostly anticoagulant and fibrinolytic [28],[29], and *L. obliqua*, whose venom mainly has procoagulant activities [3],[19]). These differences, and the suspected high inaccuracy of species identification, may explain differences in the recovery times of patients treated with LAV [13], [19].

Although accidents with venomous animals have not been the focus of much epidemiological research and accurate epidemiological data are scarce, in 2017 the World Health Organization included snakebite envenoming as a neglected tropical disease (NTD), highlighting a public health problem affecting people globally [30]. Recognizing the relevance of snakebite envenoming as a public health problem opens the door to discussing the impact of other venomous animals in vulnerable populations. In this context, it is critical to identify correctly the species present in regions where accidents occur, and to understand their toxicity, biology, and ecology.

In this study, our aims are: (1) to propose a comprehensive account of species diversity and distribution for the genus *Lonomia* over the entire Neotropical region, and (2) to interpret a newly compiled synthesis of epidemiological information in the light of this revised taxonomic assessment as a first step to improving our understanding of the areas at risk in Latin America, where surveillance and prevention strategies should be addressed locally to tackle this potentially emerging public health problem. Our results highlight the neglected diversity of venomous caterpillars and provide geographical information highly relevant to their epidemiology.

## Results

### Species delimitation and taxonomic account

Our assembled database (DS-LONO2: dx.doi.org/10.5883/DS-LONO2) comprises 1,212 records of *Lonomia* specimens. Of these records, 69% (839 records) have DNA barcode sequences and represent all species (52) and subspecies (5) listed as valid in [24], except for “*Lonomia albiguttata* Köhler, 1928”, which we consider to be just an incorrect subsequent spelling of *Lonomia albigutta* Walker, 1855 (currently placed as a junior subjective synonym of *L. obliqua*). The three most recently described species, *Lonomia cayennensis* Brechlin & Meister, 2019, *Lonomia moniqueae* Brechlin & Meister, 2019, and *Lonomia rubroguyana* Brechlin & Meister, 2019, as well as *Lonomia diabolus* Draudt, 1929, recently raised to species status by [31] from subspecies status within *L. achelous*, are also represented by DNA barcodes. Importantly, DNA barcodes are available for the holotypes of 46 of the 60 currently valid taxon names, for the lectotypes of two further species, and for paratypes of another four species, thus providing a very robust nomenclatural background (see availability of type specimen DNA barcode in DS-LONO2: dx.doi.org/10.5883/DS-LONO2) for the taxonomic account proposed here.

The analysis of genetic distances and their integration with comparisons based on morphological characters (mainly male genitalia) reveals a rather consistent taxonomic context in which the validity of most of the currently recognized species is supported by both sets of characters. Figure 2 and Supplementary Table S1 online, provide summary statistics and graphical visualization for DNA barcode variation within and among species. In total, our integrative taxonomy study leads us to recognize 53 named and seven yet unnamed species, distributed from Argentina to Mexico. We consider that these latter seven Operational Taxonomic Units (OTUs) represent undescribed species to which we here assign provisional names pending formal description. Two of these species are only known from caterpillars collected in Colombia (Amazonas) and Brazil (Mato Grosso) and are currently distinguished based solely on their very divergent DNA barcodes.

**Figure 2.**
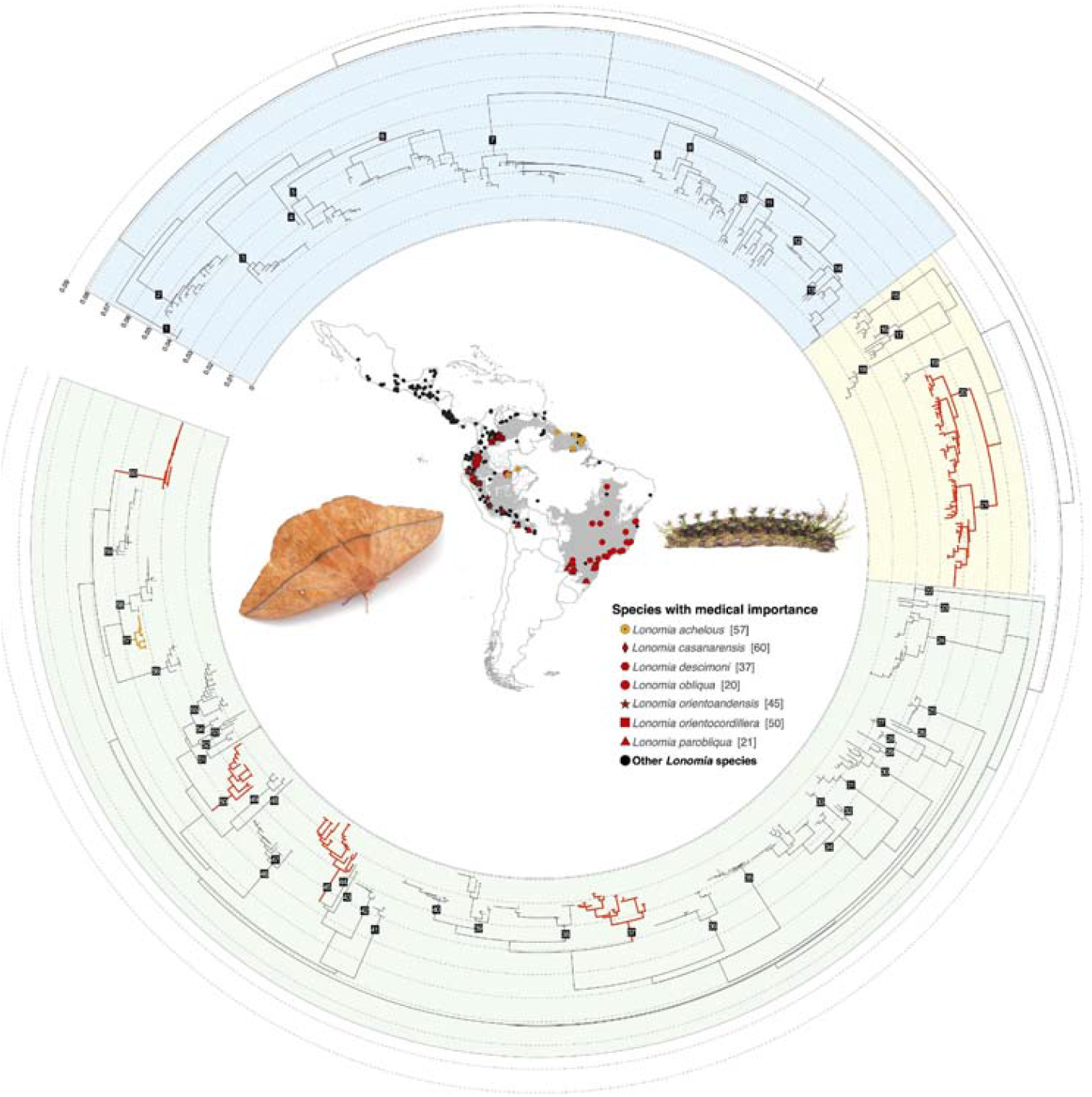
Neighbor joining tree and geographical distribution of the genus *Lonomia*. The inner map shows the distribution of the genus *Lonomia* and gray areas represent the biomes where species with medical importance have been found. Red and orange symbols in the map, and branches in the neighbor joining tree depict the species of confirmed (red) or suspected (orange) medical importance; these are listed and numbers after the species name refer to their position on the tree. Black dots depict all the other species in the genus. All species of the genus are numbered in the tree from 1 to 60 (clockwise, starting from top-left end of the tree) in reference to species as numbered in Supplementary Table S1 online. Colored background areas of the tree represent the three species-groups recognized in this work: the *electra*-group (blue), *obliqua*-group (yellow), and *achelous*-group (green). Left side picture: *Lonomia sp*. adult (live unidentified specimen from Costa Rica; courtesy of Armin Dett). Right side picture: *Lonomia camox* 6th instar caterpillar (photo by RR).

All 60 species can be identified by unique DNA barcodes, and we found no cases of identical DNA barcodes being shared by multiple species. We did observe two cases of species pairs sharing the same BIN (an automated registry of DNA barcode clusters in BOLD [32]) and further study is needed to assess these as potential cases of synonymy; these pairs are *Lonomia pastazana* Brechlin, 2017 and *Lonomia siriae* Brechlin & Meister, 2013; and *Lonomia viksinyaevi* and *Lonomia pseudobliqua* Lemaire, [1973]. In contrast, there are 15 species that split into multiple BINs (in grey in Supplementary Table S1 online), from two BINs in *Lonomia belizonensis* Brechlin, Meister & van Schayck, 2011 to eight BINs in *Lonomia sinalectra* Brechlin & van Schayck, 2015. Some of these may conceal yet unrecognized and undescribed species diversity and require further study to assess their significance.

Three subspecies were proposed by [33] and are here retained as they form distinct, though shallow, genetic clusters that are geographically segregated: *Lonomia concordia conricana* Brechlin & Meister, 2013 in northern Costa Rica and *Lonomia concordia concordia* Druce, 1886 in southern Costa Rica and Panama; *Lonomia mexilectra mexpueblensis* Brechlin & Meister, 2013 at relatively high elevation on the Volcanic Axis in Puebla and Oaxaca states of Mexico, and *Lonomia mexilectra mexilectra* Brechlin & Meister, 2011 at lower elevations along the Sierra Madre del Sur in the Mexican states of Oaxaca and Chiapas; and *Lonomia quintanarooensis quinchiapasiana* Brechlin & Meister, 2013, restricted to Chiapas state in Mexico, and *Lonomia quintanarooensis quintanaroeensis* Brechlin & Meister, 2011 distributed from southern Mexico in Quintana Roo state to Belize, Guatemala, Honduras and Guatemala (Supplementary Fig. S3 online).

Our results also lead us to propose the following taxonomic changes:

1. *Lonomia oroiana* Brechlin, Meister & Käch, 2013 syn. nov. is here synonymized with *Lonomia manabiana* Brechlin, Meister & Käch, 2013; under the Principle of the First Reviser as both species were described in the same publication. The DNA barcodes of holotype specimens of both species are nearly identical (p-distance of 0.6%), the male adult habitus and genitalia are morphologically very similar [33], and the species occupy the same biogeographical region, west of the Ecuadorian Andean cordillera. Additional morphological comparisons and DNA barcoding of a larger number of specimens would be necessary to address further the status of *L. oroiana* and so we here adopt a conservative position that limits the number of valid species within the genus.
2. *Lonomia viksinjaevi vikcuscensis* Brechlin & Meister, 2013 syn. nov. is here considered to be a junior synonym of *L. pseudobliqua*. The type localities of *vikcuscensis* and *pseudobliqua* in Cuzco and Puno departments of Peru respectively are very close (ca. 140km), and we found no differences between the male genitalia of the holotype of *L. pseudobliqua* and the illustrations of those of *L. v. vikcuscensis* in [33]. The wing pattern of the holotype of *L. pseudobliqua* is rather misleading with a yellow ground color and a double postmedial band on the forewings, which contrasts with the coloration and single postmedial band pattern observed in the holotype and two paratypes of *vikcuscensis*. However, similar intraspecific variation is known in other *Lonomia* species, for example, a Bolivian specimen of the closely related *L. viksinyaevi* Brechlin & Meister, 2011, preserved in the MNHN (sampleID: EL6149) has a single postmedial band, rather than the usual two of this species.
3. *Lonomia yucatensis* Brechlin & Meister, 2011 syn. nov. is here synonymized with *Lonomia serranoi* Lemaire (2002) a species described from El Salvador that was not considered by [34] when describing *L. yucatensis* from the Yucatan peninsula in Mexico. DNA barcode analysis of the respective holotypes places them within a single genetic cluster (BIN: AAJ2304) distributed in southern Mexico (Oaxaca, Chiapas, Yucatan), Belize, Guatemala, Honduras and El Salvador. We found no differences in male genitalia or wing patterns between specimens from the Yucatan peninsula ([34]; specimen EL6290 at MNHN) and those from other Central American regions.

Regarding the taxonomic characterization of the only two species that have been recorded so far as being of medical importance, our results clarify the identity of *L. achelous* and its distinction from the recently described and closely similar species, *Lonomia madrediosiana* Brechlin & Meister, 2011. From the habitus and genital morphology of the male neotype of *L. achelous*, as well as its geographical origin in the Guiana Shield region (Suriname), we consider that specimens collected in French Guiana and in Brazilian and Colombian Amazonia (see Supplementary Fig. S1A online), whose DNA barcodes form a cohesive genetic cluster (BOLD: ADG0261), are *L. achelous. Lonomia madrediosiana* was described from Madre de Dios department, Peru (see Supplementary Fig. S1B online), and most of its records are from the eastern side of the Andes, from southern Peru (Puno department) to northern Ecuador (Sucumbios province). However, a few specimens are also known from Ecuadorian (Orellana province) and Peruvian (Loreto department) Amazonia, which suggests that the species may extend eastward into Amazonia, where it may reach or overlap with the range of *L. achelous. L. madrediosiana* forms a separate distinct genetic cluster (BOLD: AAB4835) and DNA barcodes unequivocally discriminate the two species as circumscribed here. Morphologically, we found no clear and consistent diagnostic characters in the male genitalia to distinguish the two species. However, males of *L. madrediosiana* have a more elongated forewing with a concave external margin. In the females, the antemedial line visible in all specimens of *L. achelous* is consistently absent in the *L. madrediosiana* that we examined (*n* = 6 for both species). The apex of the forewing is also more pronounced in *L. madrediosiana* than in *L. achelous*.

The second species historically recognized as being of medical importance, *L. obliqua*, was considered by [34] to be two distinct species that he formalized through the description of the new species, *Lonomia parobliqua* Brechlin, Meister & Mielke, 2011. Our results support this distinction (see Fig. 2 and Supplementary Table 1 and 2 online), although our integrative approach combining DNA barcodes and morphology found no support for the morphological diagnostic characters proposed by [34] and reveals an overlap of the geographical ranges of both species in southeastern Brazil (see Supplementary Fig. S2 online). Our efforts to obtain a DNA barcode from the lectotype of *L. obliqua*, preserved in the Natural History Museum, London, UK (NHMUK), were in vain and we must stress that there is currently no certainty that the nomenclatural treatment of these two species is correct. Because of the medical importance of *L. obliqua*, we consider that it is currently best to follow the nomenclature proposed by [34] pending the further morphological and molecular studies that we consider necessary to assess species boundaries and the valid names to be used.

### Fieldwork campaigns

During field campaigns in Colombia (Fig 3, DS-LONO2: dx.doi.org/10.5883/DS-LONO2), we collected ten species of *Lonomia* from six different departments: *L. casanarensis* and *L. orientoandensis* were found as larvae in Casanare; in Amazonas, *Lonomia descimoni* (Lemaire 1972) was collected both as larvae and adults; adults of *Lonomia rengifoi* Brechlin & Käch, 2017 and *L. achelous* were also collected in Amazonas, as well as caterpillars of an unidentified species (provisionally named as *Lonomia* CGR01). In Risaralda, light-trapping yielded large numbers of adult *Lonomia columbiana* Lemaire, 1972. In Meta, we identified *L. descimoni* by DNA barcoding caterpillars involved in an accident that occurred after contact of the victim with a colony of caterpillars on the trunk of a “Guamo” tree (*Inga* sp.). We also obtained DNA barcodes from different caterpillars at the exact same site that belong to *Lonomia orientocordillera* Brechlin *et al*., 2013. These findings in Meta are the first formal confirmation that species other than *L. achelous* and *L. obliqua* are involved in human accidents. In Guainía (2018), an adult female belonging to *Lonomia cayennensis* Brechlin & Meister, 2019 was collected in a field peripheral to the city of Inirida. Finally, in 2019, we collected four specimens of *Lonomia minca* Brechlin, 2017 in Magdalena department. We found no *Lonomia* caterpillars during our campaigns in French Guiana, but light-trapping yielded material from six different species: *L. belizonensis, Lonomia beneluzi* Lemaire, 2002, *L. cayennensis, Lonomia camox* Lemaire, 1972, *L. diabolus*, and *Lonomia rubroguyana* Brechlin & Meister, 2019 (Supplementary Fig. S1A and S1B online).

**Figure 3.**
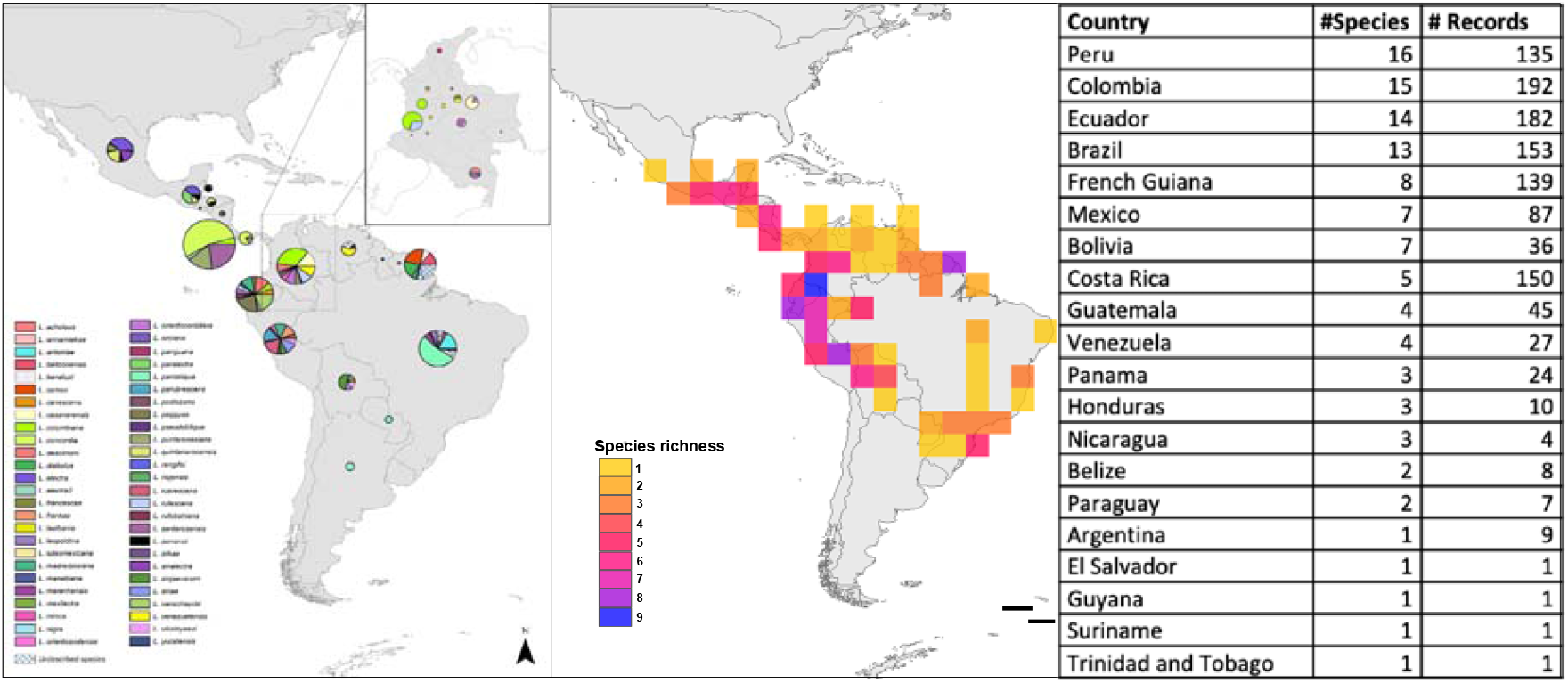
*Lonomia* species richness. Left side map illustrates species richness in Latin America by country, and in the inset a map of Colombia by department. Centre map shows species richness in Latin America at a resolution of 400 km^2^ grid size, and the right-side table shows the number of *Lonomia* species recorded in each country, and the number of collection records. The list of countries is sorted in decreasing order of number of species.

### Spatial distribution

Overall, we successfully assigned geographic coordinates to 1,019 records, representing 585 localities for the species recognized here. The spatial distribution of the genus *Lonomia* extends across the entire Neotropical region in Central and South America, from Mexico to northern Argentina, although it is absent from Chile and Uruguay (Fig. 2).

Species richness was greatest in Peru, with 19 species recorded, followed by Colombia with 15 species (including one undescribed), Ecuador with 14 species, and Brazil with 13 species (including five undescribed). Regarding the number of collection localities, Colombia is the best sampled country (*n* = 183), followed by Ecuador (n=169), Costa Rica (n=137), French Guiana (*n* = 115), Brazil (*n* = 113), and Peru (*n* = 93) (Fig.3, Supplementary Fig. S1A – S3 online).

The species with the broadest ranges (measured by the number of countries inhabited) are *L. achelous* (Brazil, Colombia, French Guiana, Guyana, Suriname) (Fig. 3, Supplementary Fig. S1A online;) and *L. parobliqua* (Argentina, Bolivia, Brazil, French Guiana, Paraguay) (Fig.3, Supplementary Fig. S2 online); in South-America, and *L. quintanarooensis* (Belize, Guatemala, Honduras, Mexico, Nicaragua) (Fig. 3, Supplementary Fig. S3 online); and *L. serranoi*, (Belize, El Salvador, Guatemala, Honduras, Mexico) in Central America (Fig. 3, Supplementary Fig. S1B online).

### Coincidence with areas of epidemiological relevance

Our survey of the available epidemiological information produced data for six countries in South America: Argentina, Brazil, Colombia, Peru, Paraguay and Venezuela, as well as French Guiana, with records of hemorrhagic syndromes after accidental contact with caterpillars of *Lonomia* (Supplementary Table S2 online). In Argentina, most cases have been recorded in Misiones, where eight cases from seven localities were documented between 2010 and 2015. Five of these involved children under the age of 17 years, one of them fatal despite the use of antivenom (Supplementary Table S2 online). In Argentina and Paraguay, only one species has been reported to be involved in accidents, historically named as *L. obliqua*. However, the Argentinian populations of *L. obliqua* were considered by [34] to be a distinct species, *L. parobliqua*, and DNA barcodes support the conclusion that this species is the only one present in this country. A DNA barcode sequence obtained from a specimen involved in an accident in Misiones was a match for *L. parobliqua* (Maria Elisa Peichoto, personal communication, 21 Sep. 2017). Although Corrientes province in Argentina is considered an area of epidemiological importance [35], we could not locate any specimen collected in that province.

In Brazil, 1,930 accidents involving *Lonomia* caterpillars were recorded between 2001 and 2006, 70% of which were in the southern and 20% in the southeastern regions of the country (Fig 4). From 2007 to 2017, 27,263 accidents with caterpillars (not only *Lonomia*) were recorded, 42% in the southern, 39% in the southeastern, and 10% in the northeastern parts of Brazil. Out of these, 33 were fatal envenomation accidents, 54% occurring in the southern region of the country.

**Figure 4.**
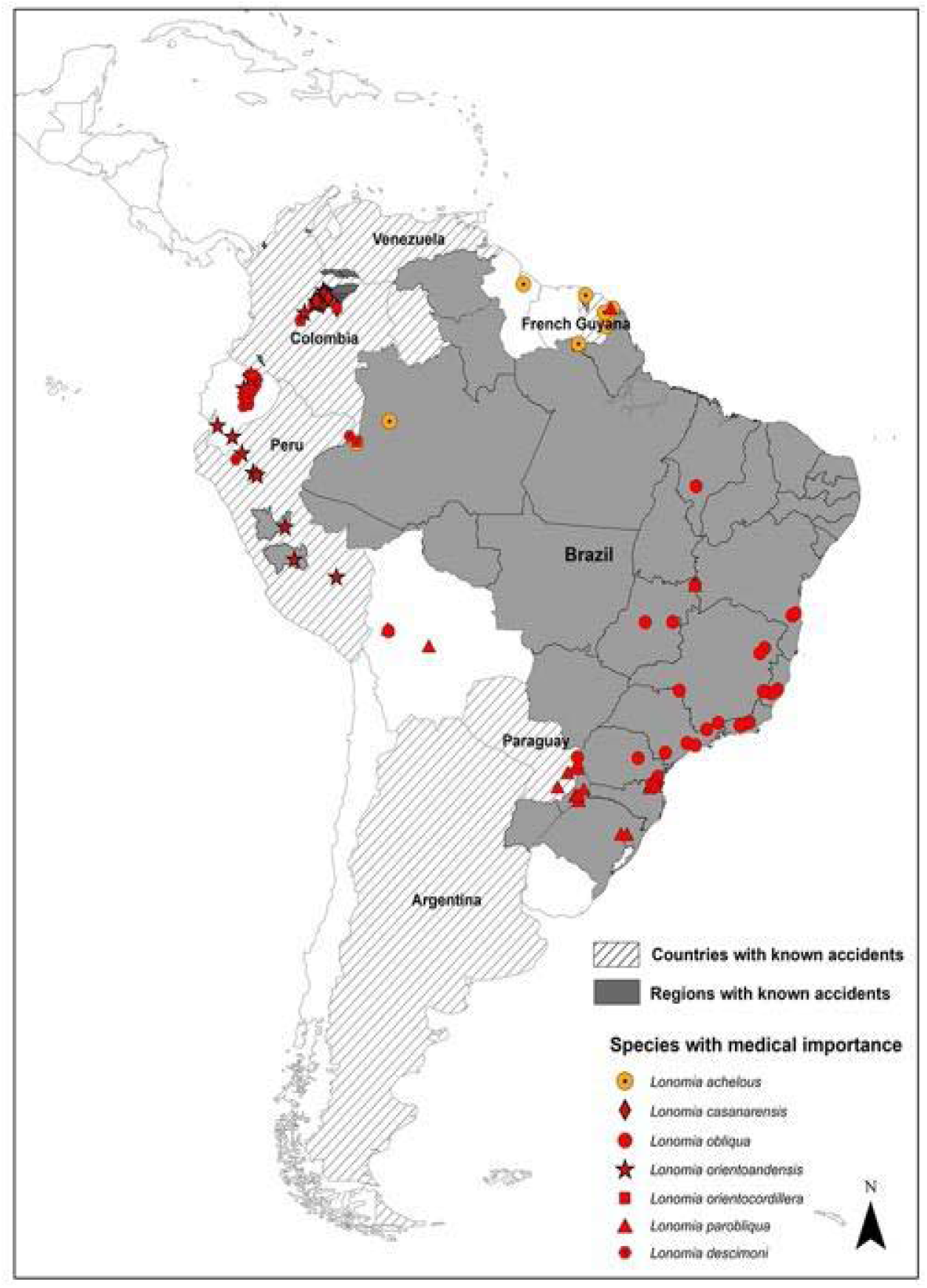
*Lonomia* accidents. The map shows the administrative unit level for which information about accidents with *Lonomia* caterpillars is available. Dashed lines: country level, solid gray: states or municipalities. Red and orange symbols represent the distributions of the species of confirmed (red) or suspected (orange) medical importance in South America.

In all states where accidents have been recorded (except Rio Grande do Sul) *L. obliqua* is present. However, other species are also present in this area, such as *Lonomia antoniae* Brechlin & Meister, 2015 in Minas Gerais, Paraná, Santa Catarina and Sao Paulo (Supplementary Fig. S2 online), *Lonomia leopoldina* in Bahia, Rio de Janeiro and Santa Catarina (Supplementary Fig. S2 online), *Lonomia maranhensis* in Maranhão (Supplementary Fig. S1A online), *L. parobliqua* in Goias, Paraná, Rio Grande do Sul and Santa Catarina (Supplementary Fig. S2 online), *Lonomia rufobahiana* Brechlin & Meister, 2013 in Bahia (Supplementary Fig. S2 online) and the undescribed species, *Lonomia* RRBR01 in Paraná and Santa Catarina (Supplementary Fig. S2 online). The state with the greatest number of recorded deadly accidents is Rio Grande do Sul where *L. parobliqua* is present, but from where no records of *L. obliqua* exists.

In French Guiana, three cases of envenomation by *Lonomia* have been reported in the last 25 years. The most recent occurred in 2017 in Saint Laurent du Maroni and was successfully treated with LAV [19]. The species responsible could not be identified by the authors but they suspected *L. achelous* based on the two previous accidents reported by [18]. Our results reveal that there are at least four *Lonomia* species present in the area where this recent accident occurred (i.e., western French Guiana), two of which have only recently been described: *L. achelous, L. rubroguyana, L. cayennensis* and *L. camox* (Supplementary Fig. S1A and Fig. S2 online).

Peru has the highest species richness, and accidents have been recorded in Junín in 2006, and in Huánuco in 2010 and 2017 when antivenom was used (Supplementary Table S2 online). The species reported to be present here are: (1) in Junín, *L. orientoandensis, L. madrediosiana, Lonomia silkae* Brechlin & Meister, 2013 and *Lonomia frankae* Meister, Naumann, Brosch & Wenczel, 2005; (Supplementary Fig. S1A and Fig. S1B online) (2); and in Huánuco, *Lonomia parubrescens* Brechlin & Meister 2011, *L. orientoandensis* and *Lonomia panguana* Brechlin, 2017, (Supplementary Fig. S1A online).

In Venezuela, from 1960 to 1967, five accidents of hemorrhagic syndrome caused by contact with venomous caterpillars were recorded in Bolivar state (Arocha Piñango, 1967). *Lonomia achelous* was, as usual, recorded as the causal agent (Arocha Pinango, 1967) but only one species, *Lonomia moniqueae* Brechlin & Meister, 2019, has been collected in that state (Supplementary Fig. S1A online). In 1996, caterpillar colonies were collected from *Tapirira guinanensis* tree trunks in Anzoategui and Bolivar states [36] but we have no *Lonomia* collection records from Anzoategui in our database (Fig. 4).

In Colombia, the first reported envenomation accident by *Lonomia* caterpillars was fatal for its victim. It occurred in 2000 in San Nicolás, Nunchía, department Casanare [13]. Although a female moth reared from the caterpillars that caused the accident was identified at the time as *L. achelous*, its DNA barcode now identifies it as *L. casanarensis* (SampleID: CGR_Lon34). From Meta department, we were able to identify through DNA barcoding specimens provided for identification purposes by the Secretaría de Salud del Meta that were linked to three accidents, which proved to be *L. descimoni* (CGR_Lon84 and CGR_Lon90; in Cubarral, 2017), *L. casanarensis* (CGR_Lon91, in Acacías, 2017), and *L. orientocordillera* (CGR_Lon85 and CGR_Lon86, in Cubarral, 2017). The National Institute of Health also provided information on six accidents in 2017, one in Casanare (Orocué), four in Meta (one from Acacías, one from Cubarral and two without information) and one in Arauca (Saravena). In all cases the victims attended hospital, and at least five received LAV provided by Butantan as treatment. The *Lonomia* species collected in Casanare, where most accidents have been recorded are *L. casanarensis, L. orientoandensis* and *L. orientocordillera* (Supplementary Fig.S1A online). In Meta, these are also present together with *L. descimoni* (Supplementary Fig. S1A online). In Arauca department, although accidents have been reported, we have no formal records of *Lonomia* and no specimens are available that would permit further investigation of their identification (Fig.4).

## Discussion

For the first time, integrated information at a continental scale on *Lonomia* venomous caterpillars has been compiled and made accessible to expose a poorly known but potentially emerging public health problem. Through an integrative approach combining genetic, morphological and geographical data, we propose a comprehensive appraisal of the diversity of this genus that confirms the presence of 60 distinct species in America, in striking contrast to the 11 species recognized by Lemaire [23]. Our results validate most of the scattered taxonomic work achieved in the past 20 years and significantly impact and clarify our understanding of the identities of the two species traditionally considered to be of medical importance, *L. achelous* and *L. obliqua*. Regarding *L. obliqua* in Brazil, where it is the only documented species related to human envenoming, it is important to establish its identity and to define both its spatial distribution and the epidemiological importance of the two overlapping species, *L. obliqua* and *L. parobliqua*. In addition, we have firmly established the identity of several other species involved in accidents with humans: *L. casanarensis, L. orientoandensis, L. orientocordillera, L. parobliqua* and *L. descimoni*. From distributional information, there is now compelling evidence that many reported cases of human envenomation have been erroneously attributed to *L. obliqua* and *L. achelous*, and that in addition to the aforementioned species, it is highly likely that the caterpillars of other species (e.g., *L. madrediosiana, L. moniqueae, L. antoniae*) can also cause severe envenomation in humans. In Colombia alone, three different species were reported as involved in envenomation syndromes: *L. descimoni, L. orientocordillera* and *L. casanarensis*. A fourth, *L. orientoandensis*, has been shown to induce hemorrhagic syndrome in rat [6] and should be considered as a potential health hazard for humans as this widely distributed species occurs in areas where accidents have been notified in both Colombia and Peru. Importantly, species involved in accidents can be endemic with narrow distributions, such as *L. casanarensis* in the Colombian Llanos region, which emphasizes the need to gain an accurate knowledge of species diversity and distribution to efficiently address envenomation risks and define possible treatments.

The use of barcodes for the identification of *Lonomia* species causing human envenoming is of great help, providing a rapid and reliable tool for decision making and should be included as a routine part of response strategies.

With the information provided in this work, health authorities can increase surveillance in areas at risk and design adequate response strategies encompassing the enormous species diversity and wide distribution of the genus. Although accidents with venomous caterpillars are not (yet) considered neglected tropical diseases, their epidemiological patterns match such, in that they affect impoverished communities and face strong challenges regarding diagnostics and treatment [36].

Interestingly, despite the wide distribution of the genus *Lonomia*, accidents are only known from South America, whereas in Mexico and Central America, to our knowledge, no accidents have been reported despite the presence of 11 species. Being able to quantify the problem of accidents with *Lonomia* in South America is of great importance. We hypothesize that there may be variation in toxicity accompanying the enormous diversity of the genus, since in Central America, it has been anecdotally documented that contact with *Lonomia electra* Druce, 1886 does not trigger hemotoxic activity although it does trigger urticaria upon skin contact with the venomous spines (https://entomologytoday.org/2017/03/23/up-close-and-personal-with-venomous-moths/lonomia-electra-skin-irritation/, [37]). Although the widely used treatment, LAV, produced against *L. obliqua* has proven to be effective for at least two other *Lonomia* species [6], it would be important to further explore its efficacy against other species. For example, until 2020, accidents in Colombia with species of *Lonomia* other than *L. achelous* were successfully treated with the antivenom produced by the Butantan Institute against *L. obliqua* and in French Guiana the maximum recommended dose (10 vials) of the same antivenom was used [38],[19]. Thus, there is an opportunity for an integrated initiative leading to the design of treatment protocols in case of accidents with *Lonomia*, where species identification will be a must, and perhaps the production of a polyvalent treatment that can be used in different countries, as has been proposed for snakebite [39], [40].

Due to loss of natural habitat and possibly attraction to the lights of human habitations, *Lonomia* species originally occurring in forested areas are shifting their distributions into urban settings in their search for host plants, which are often found near rural houses, or used as fences in oil palm plantations, and thereby become a potential occupational risk [13]. In the future, environmental changes, particularly those related to increasing deforestation rates in tropical America, will make vulnerable human populations more prone to suffering accidents with venomous animals. For ectothermic animals such as snakes, it has been shown that changes in temperature and rainfall regimes across time affect the incidence of snakebites [41], and that rainfall seasonality seems to modulate snakebite incidence [42]. Improving our understanding of the ecological patterns, seasonal distribution, shifts with environmental changes and epidemiology of accidents with venomous animals will be the necessary focus of forthcoming research.

In conclusion, our work makes an important contribution to the field of venomous animals and particularly insects with medical importance, as well as highlighting the relevance of closing biodiversity knowledge gaps and increase sampling efforts in groups such as moths where information is particularly scarce in the most biodiverse areas of the planet.

## Material and methods

### Taxonomic account

To address the diversity of species within the genus *Lonomia* adequately and comprehensively, we used an integrative approach combining comparative morphology, DNA barcoding and biogeography, targeting all species and subspecies listed in the most recent available checklist for bombycoid moths [24], plus three additional species described subsequently [43] and another recently raised to species status by [31]. The material used in these analyses included: (1) specimens sampled from multiple public and private collections in the context of the DNA barcoding campaigns for Lepidoptera [44]; (2) newly collected specimens from targeted field work carried out by several of the authors in Colombia and in French Guiana; and (3) specimens in the Lemaire collection (CLC) at the Muséum national d’Histoire naturelle (MNHN) in Paris - one of the largest and most comprehensive collections of *Lonomia* moths in the world. All records used in this work can be accessed in the publicly available BOLD dataset DS-LONO2: dx.doi.org/10.5883/DS-LONO2). When this work began, there were 12 taxa recognized in the CLC corresponding to the 11 species and one subspecies recognized in the revision of subfamily Hemileucinae by [23]. We databased records for all specimens in this collection and organized them based on their general habitus (wing pattern) and their geographical origin to sort specimens as candidate representatives of the many species described after 2002. Then we selected individuals of both sexes within each of the sorted sets of specimens and processed them through DNA barcoding (see below) to compare them with DNA barcodes available from other sources (including type specimens of most of the recently described taxa). Dissections of male genitalia were also carried out following a standard protocol [45]. These provided an additional set of characters for species identification and delineation, to allow direct comparisons with the illustrations of these structures published in the original descriptions of recognized species.

### Fieldwork campaigns

To identify which species were present in areas of Colombia where accidents have been reported, fieldwork was performed between 2012 and 2019 to capture caterpillars and adults in different regions of this country (DS-LONO2: dx.doi.org/10.5883/DS-LONO2). When caterpillar colonies were found, all individuals from each colony were collected and a group of up to ten individuals was reared to obtain adults [46], while most of the remaining caterpillars were used for venom extraction following the methods of [6]. At least two caterpillars from each colony were preserved in 70 % ethanol and tissue samples were obtained to confirm species identity through DNA barcoding (as detailed below) when rearing was not possible. We also collected adult moths with a light trap using a 125W mercury vapor bulb powered by a small portable generator, with a 2×3m white sheet used as a reflector. Trapping was performed from dusk to dawn (18:30 to 6:30). Moths were collected, injected with ammonia, then stored dried in labeled paper envelopes for later identification using genital morphology and DNA barcoding. Light-trapping using the same method was also carried out in French Guiana during three field trips in the Nouragues nature reserve (2011), in the French Agricultural Research Centre for International Development (CIRAD) research station of Paracou (2019) and in the Kaw mountains (2020) (DS-LONO2: dx.doi.org/10.5883/DS-LONO2).

### Laboratory procedures and DNA barcode analyses

For newly collected specimens (both adults and caterpillars), DNA was extracted from one or two legs following the standard protocol described in [47], using available kits (Promega Corp., Madison, WI, USA; QIAGEN DNeasy Tissue and Kit gDNA Mini Tissue ChargeSwitch® Invitrogen Corp., Carlsbad, California). After extracting DNA, the pair of primers LepF1/LepR1 was used to amplify a 658 bp fragment of the COI gene [48],[44]. PCR conditions for a volume of 50 μL were: 4.0 mM MgCl2, 0.2 μM of each primer, 0.2 mM dNTPs, 1× Buffer and 1.25 U of Taq DNA polymerase. After visualization by electrophoresis, successfully amplified products were purified and sequenced in both directions using the same primers as used for the PCR. Sanger sequencing was carried out using the ABI 3500 Genetic Analyzer sequencer at the Sequencing Laboratory of Universidad de los Andes. Sequence alignment, editing and assembly of contigs was performed using Sequencer version 6.1.0 (Gene Code Corporation, Ann Arbor, U.S, Michigan).

In addition, *Lonomia* specimens from the Lemaire collection at the Muséum national d’Histoire naturelle (MNHN, Paris, France) were processed in the molecular biology laboratory platform (‘Service de Systématique Moléculaire’ – SSM) of the MNHN. DNA extraction was carried out using Macherey-Nagel Nucleospin® 96 tissue kits following manufacturer’s protocol using either a semi-automated procedure with an Eppendorf Liquid Handling WorkstationepMotion® 7075 VAC, or a manual approach through successive centrifugations. The primer pair LepF1/LepR1 was used in a first round of amplification; PCR products were then checked on a 2% agarose gel and successfully amplified DNA templates were sent for Sanger sequencing on an ABI 3730XL sequencer at Eurofins MWG Operon sequencing facilities (Ebersberg, Germany). Failed samples (likely due to DNA degradation) were re-processed using the primer combinations LepF1/MLepR1 and MLepF1/LepR1 targeting two overlapping fragments of 307bp and 407bp, respectively [44]. PCR products were checked on a 2% agarose gel and successful samples were sequenced on an Ion Torrent PGM platform at SSM using the protocol described in [49]. All the reads obtained were analyzed and assembled with the software Geneious V11.0.4 [50].

Other DNA barcodes of relevant *Lonomia* samples were obtained from the results of the global DNA barcoding campaign for Saturniid moths and the Lepidoptera inventory program in Area de Conservación Guanacaste (ACG) in northwestern Costa Rica [51]. These were generated at the Canadian Centre for DNA Barcoding (CCDB, Centre for Biodiversity Genomics, University of Guelph, Ontario, Canada) following standard protocols [52].

All DNA barcode records used in this study, including both specimen (e.g., voucher repository, identification, collecting data, GPS coordinates) and sequence data (e.g., electropherograms, DNA sequence, GenBank accession numbers), are integrated into the BOLD dataset DS-LONO2 (dx.doi.org/10.5883/DS-LONO2) within the Barcode of Life Datasystems (BOLD, www.boldsystems.org;[53]).

We used BOLD analytical tools to compute genetic distances (using default parameters, BOLD sequence alignment option and uncorrected p-distances), and to represent them in the form of Neighbor Joining (NJ) tree. For visualization purposes, the BOLD NJ tree was imported as newick format and edited into iToL v4 [54]. Barcode Index Numbers (BINs) – i.e., molecular taxonomic units automatically assigned by BOLD [32] – are used for convenience in our presentation and discussion of the results; they are reported here as computed on BOLD on July 15th, 2022.

### Spatial distribution and coincidence with areas of epidemiological relevance

Once our database was assembled, we georeferenced those records that lacked coordinates using the locality reported on specimen labels. We used Google Earth and various online gazetteers, atlases, and traveler blogs to retrieve geographical coordinates as precisely as possible.

Epidemiological information was gathered from published case reports or through Google® searches using the terms “*Lonomia* accidents” in both English and Spanish. Information from Brazil was obtained from the National Information System SINAN (http://portalsinan.saude.gov.br/dados-epidemiologicos-sinan) and information from Colombia was provided by the National Institute of Health. Municipality boundaries were gathered from GADM data (version 3.6, https://gadm.org/about.html). Spatial analyses were performed using ArcGis 10.7 (Licensed to Uniandes).

## Data Availability

The dataset generated and analyzed in this study is available in the Bold repository, DS-LONO2: dx.doi.org/10.5883/DS-LONO2.

## Acknowledgements

We wish to give special thanks to our local collaborators at the field sites in Tanimboca Natural Reserve and Secretaría de Salud (Amazonas), Senderos del Canajagüa (Meta) and La Victoria, Caoba Natural Reserve, and Eudes Baca and his family (Magdalena), who supported this research with their motivation of contributing to scientific knowledge, aware that ultimately it can be translated into benefits for the people living in areas at risk. In the same spirit, we want to thank: Gabriel Colorado, Maria Elisa Peichoto and Pedro Sarmiento, who contributed specimens for barcode identification; the team at CIMPAT and the Department of Biological Sciences, who were always prompt to support sample processing; Giovanni Randazzo, for imaging documentation during fieldwork and Alessandro Giusti (NHMUK) for images of the holotype of *L. pseudobliqua* (including its dissected genitalia). Silvia Restrepo and Vicerrectoría de Investigaciones Uniandes, who were essential for funding this research; and The National Institute of Health INS, Colombia, who provided information on accidents recorded in the country in 2017. We are also grateful to the Canadian Centre for DNA barcoding and Centre for Biodiversity Genomics at University of Guelph (Ontario, Canada), as well as participants to the Saturniidae DNA barcoding campaign for their contribution to assemble the DNA barcode library of *Lonomia* moths. Funding: Vicerrectoria de Investigaciones, and Facultad de Ciencias INV-2020-105-2030, Universidad de Los Andes (Granted to CG). Pontificia Universidad Javeriana, Facultad de Estudios Ambientales y Rurales, PPTA 7996 (Granted to AR A-S). Humboldt Institute through the collaboration agreement 16-220. French National Research Agency (ANR) SPHINX grant no. ANR-16-CE02-0011-01 (to RR, TD) and French Foundation of Research on Biodiversity (FRB; www.fondationbiodiversite.fr) and CESAB synthesis centre to ACTIAS project (to RR, LBM, IJK, TD).

## Author contributions

Conceptualization (CG, ISSM, ARAS, RHPM, IJK & RR), Data providing (CG, JDD, DMTV, NR, LSB, RHPM, TD, IJK & RR), Data curation (CG, JDD, DMTV, LBM, MA, MAG, IJK & RR), Molecular analysis (JDD,DMT,CL, MAG, ET & DG), Visualization (CG, JDD, LBM & RR), Writing original draft (CG, JD,DMTV, LBM, IJK & RR). All authors reviewed and approved the manuscript.

## Data availability

Detailed data are available in DS-LONO2: dx.doi.org/10.5883/DS-LONO2.

## Ethics declarations

The author declare no competing interests.

