## Supplementary figures and images for "Deadly and venomous *Lonomia* caterpillars are more than the two usual suspects"

### Supplementary Fig. S1A

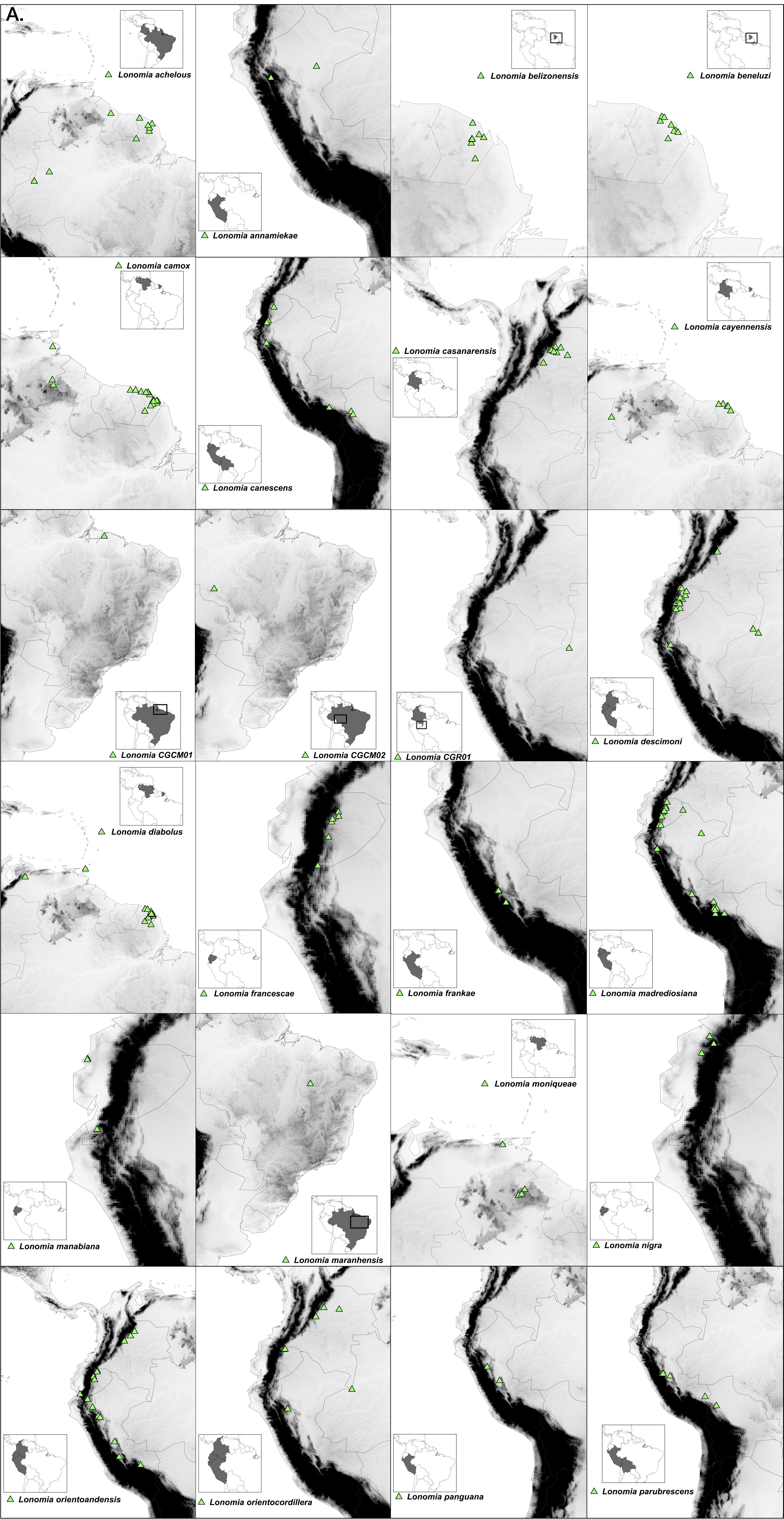
