## Supplementary Fig. S1B for "Deadly and venomous *Lonomia* caterpillars are more than the two usual suspects"

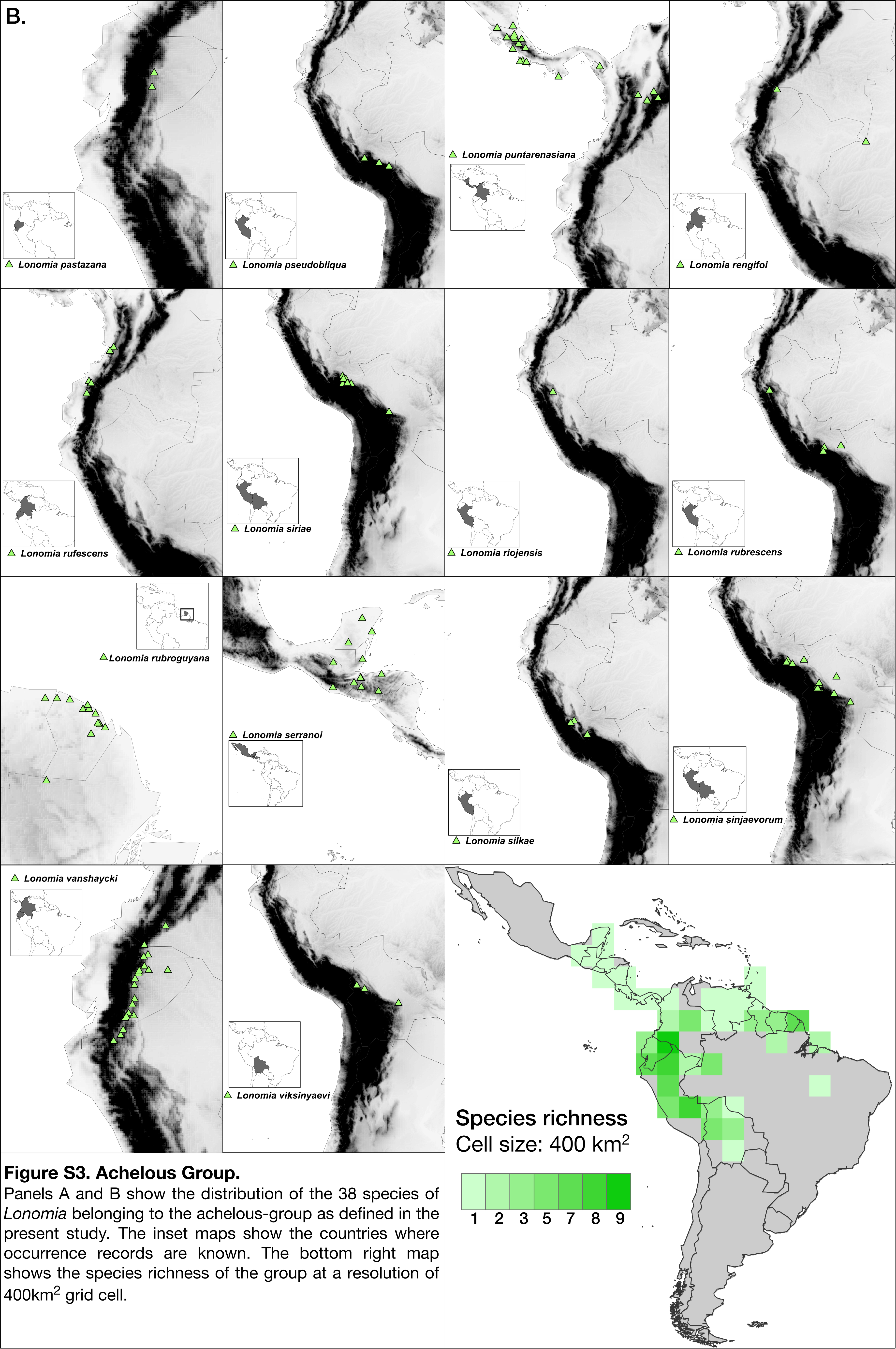


**Figure S3. Achelous Group.**
Panels A and B show the distribution of the 38 species of *Lonomia* belonging to the achelous-group as defined in the present study. The inset maps show the countries where occurrence records are known. The bottom right map shows the species richness of the group at a resolution of 400km$^2$ grid cell.
