## Supplementary Fig. S2 for "Deadly and venomous *Lonomia* caterpillars are more than the two usual suspects"

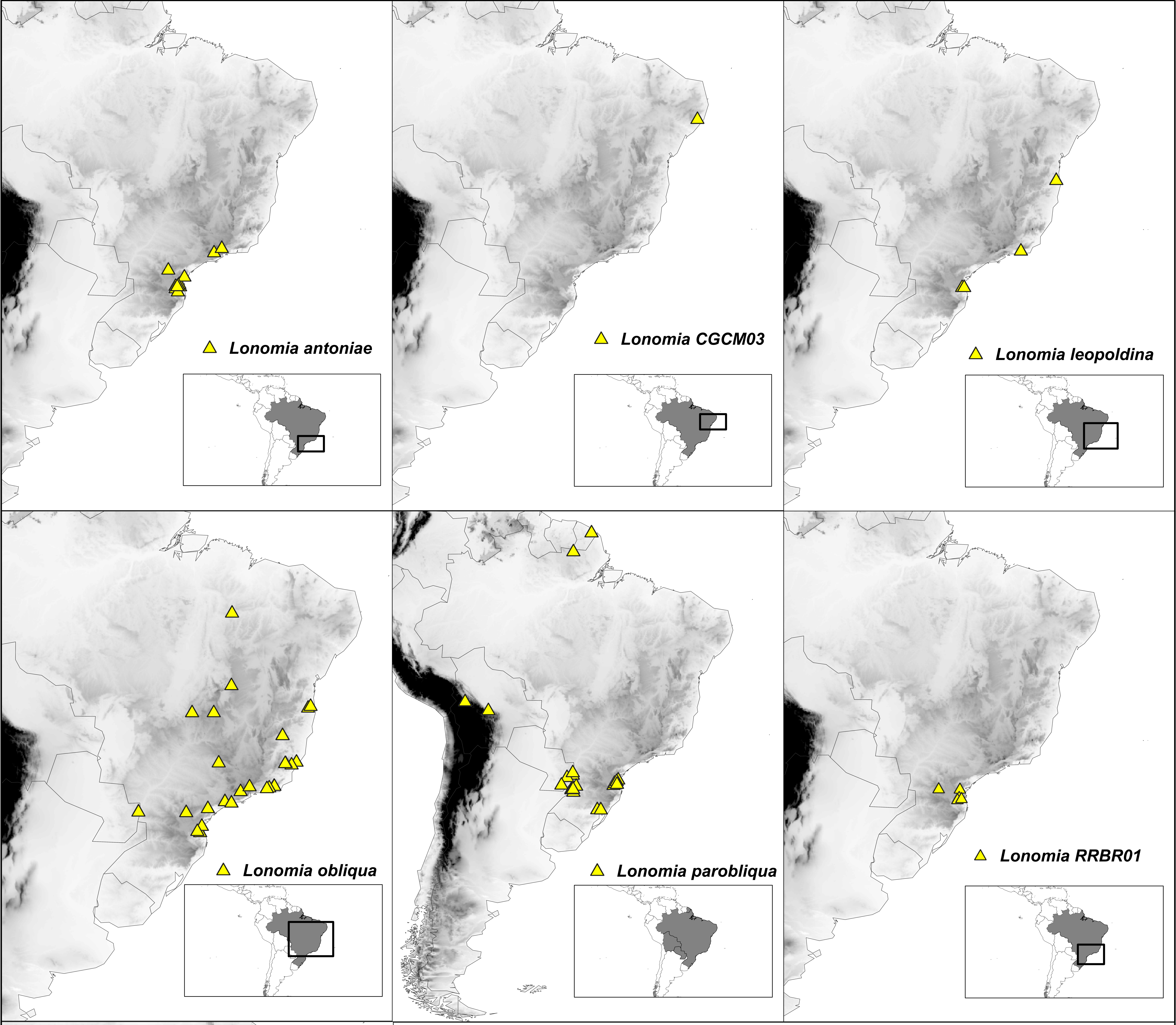


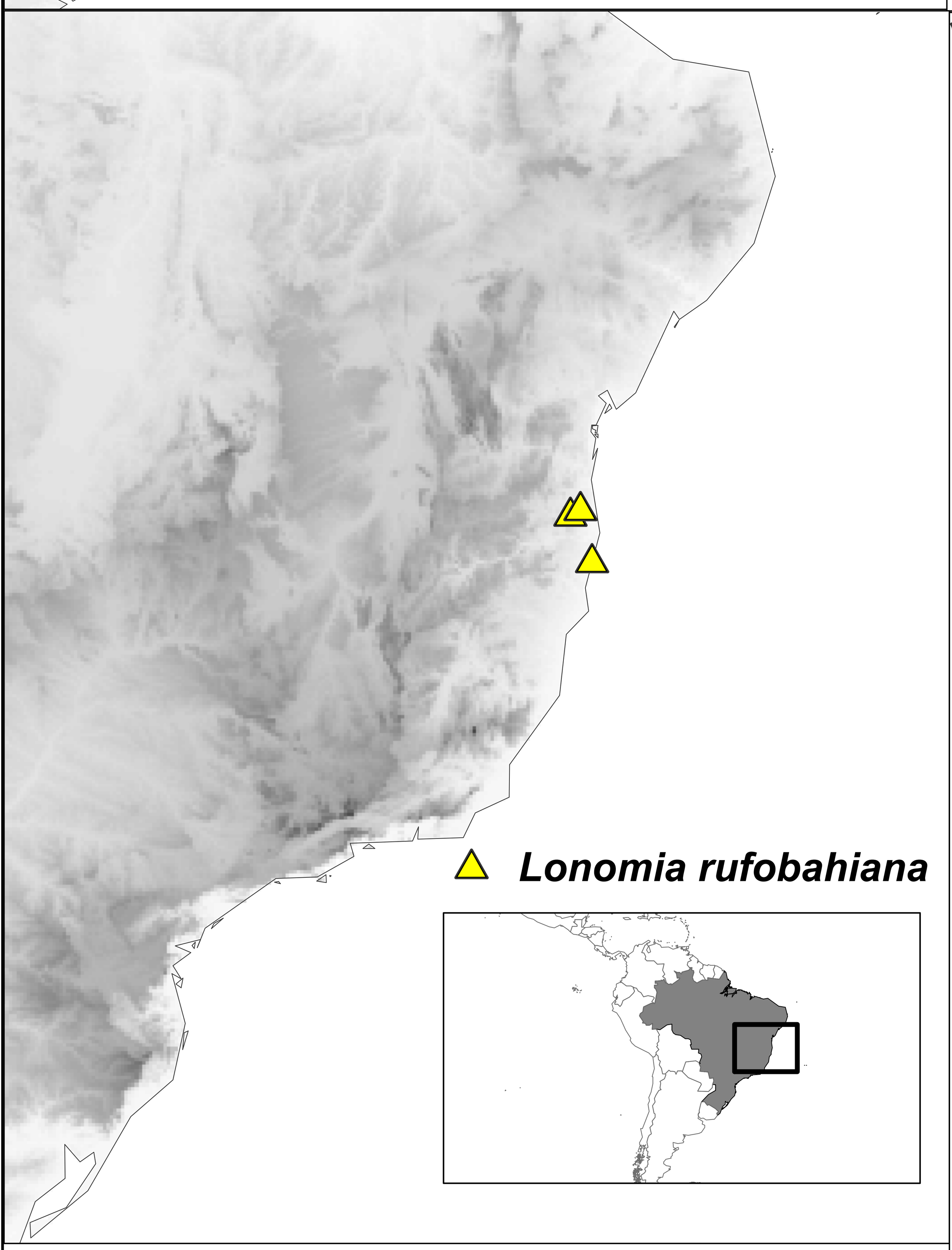


**Supplementary Fig. S2. Obliqua Group**
This panel shows the distribution of each one of the seven (7) *Lonomia* species belonging to the Obliqua group. The inset maps show the countries in South and/or Central America where the occurrence records have been recorded and a square is present in the inset maps in case a more precise information of the location within the country is needed. The map in the bottom right corner shows the distribution of the species richness of this group at a resolution of 400km$^2$ grid cell.

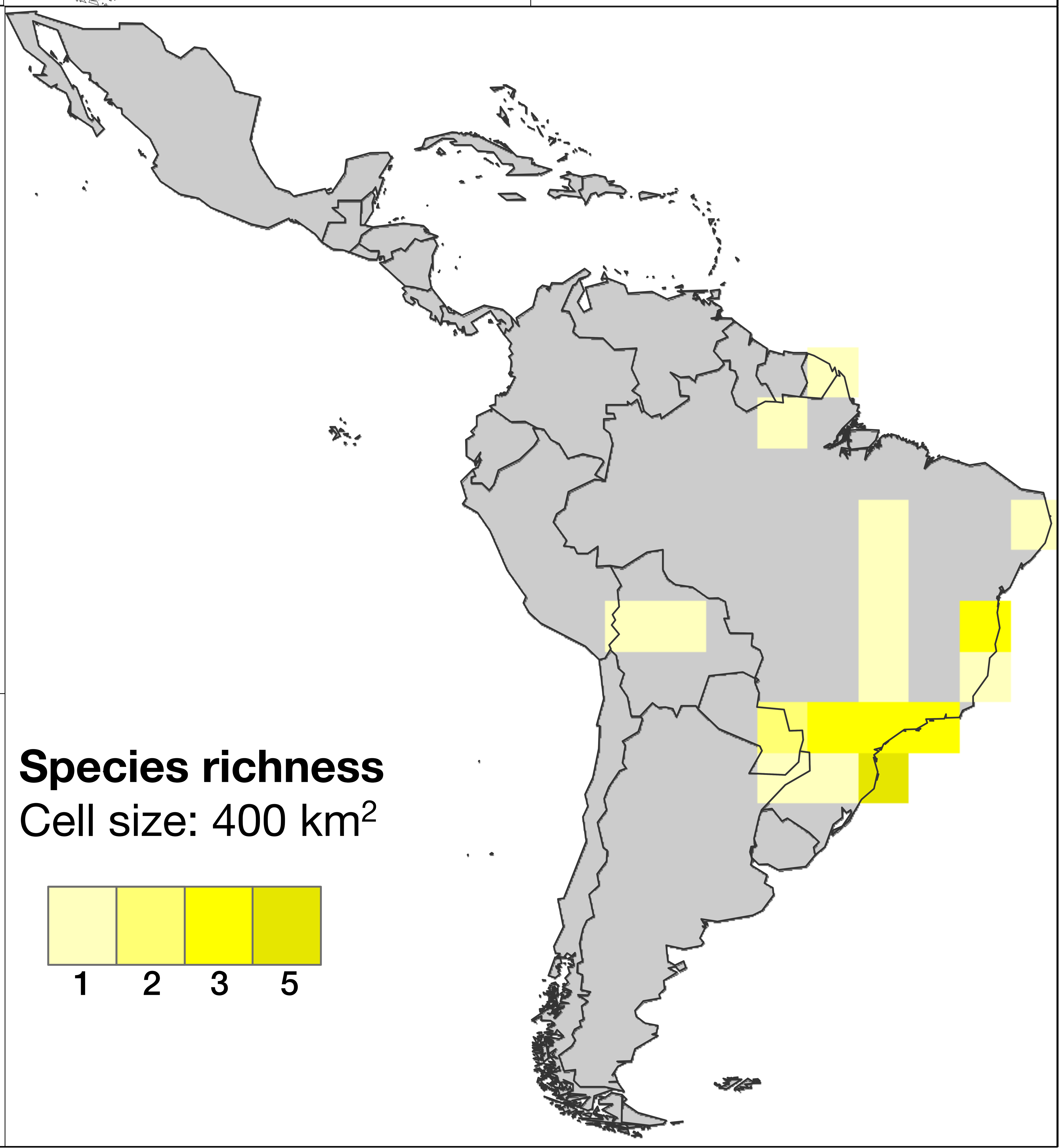
