## Supplementary Fig. S3 for "Deadly and venomous *Lonomia* caterpillars are more than the two usual suspects"

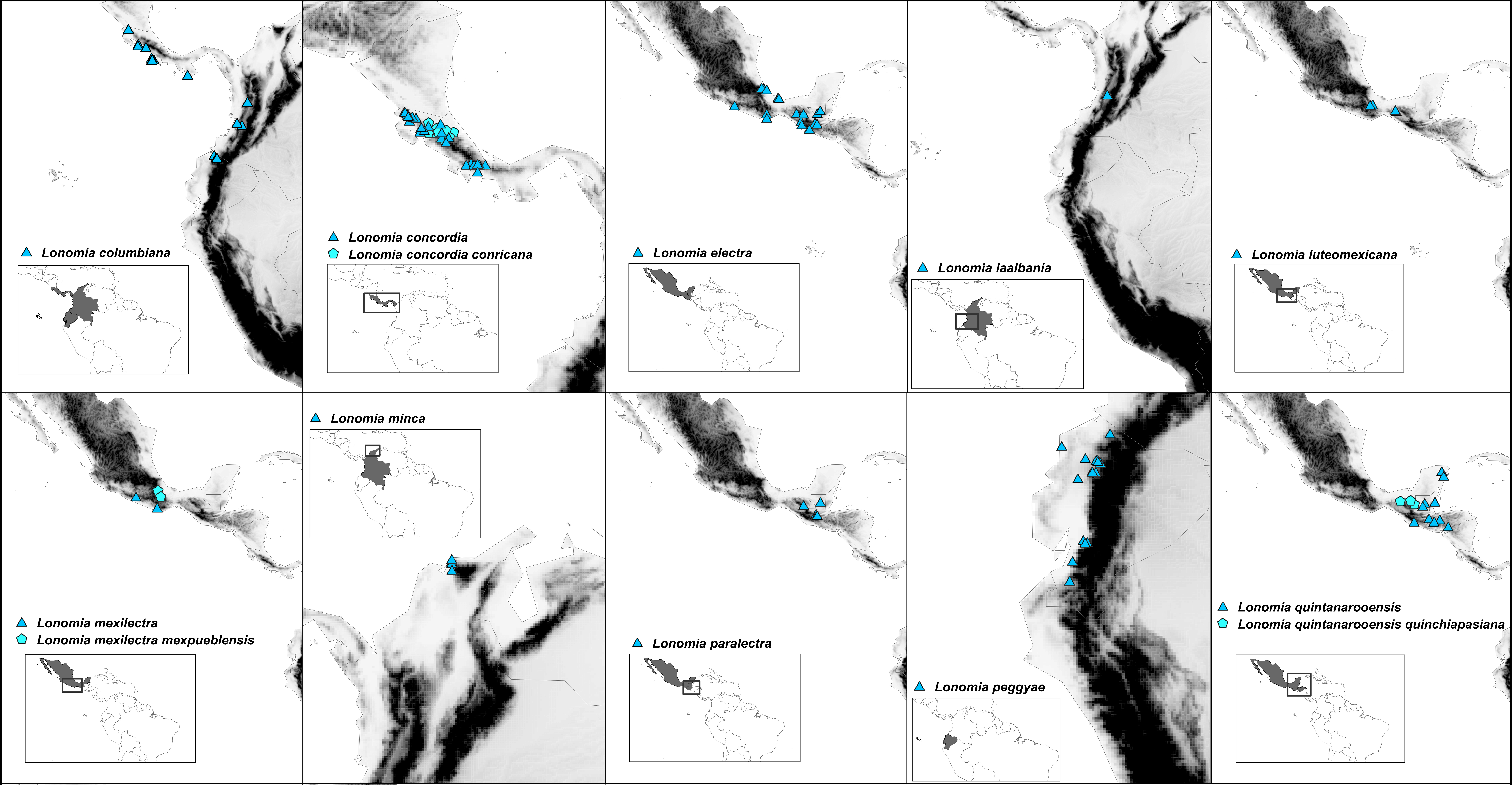


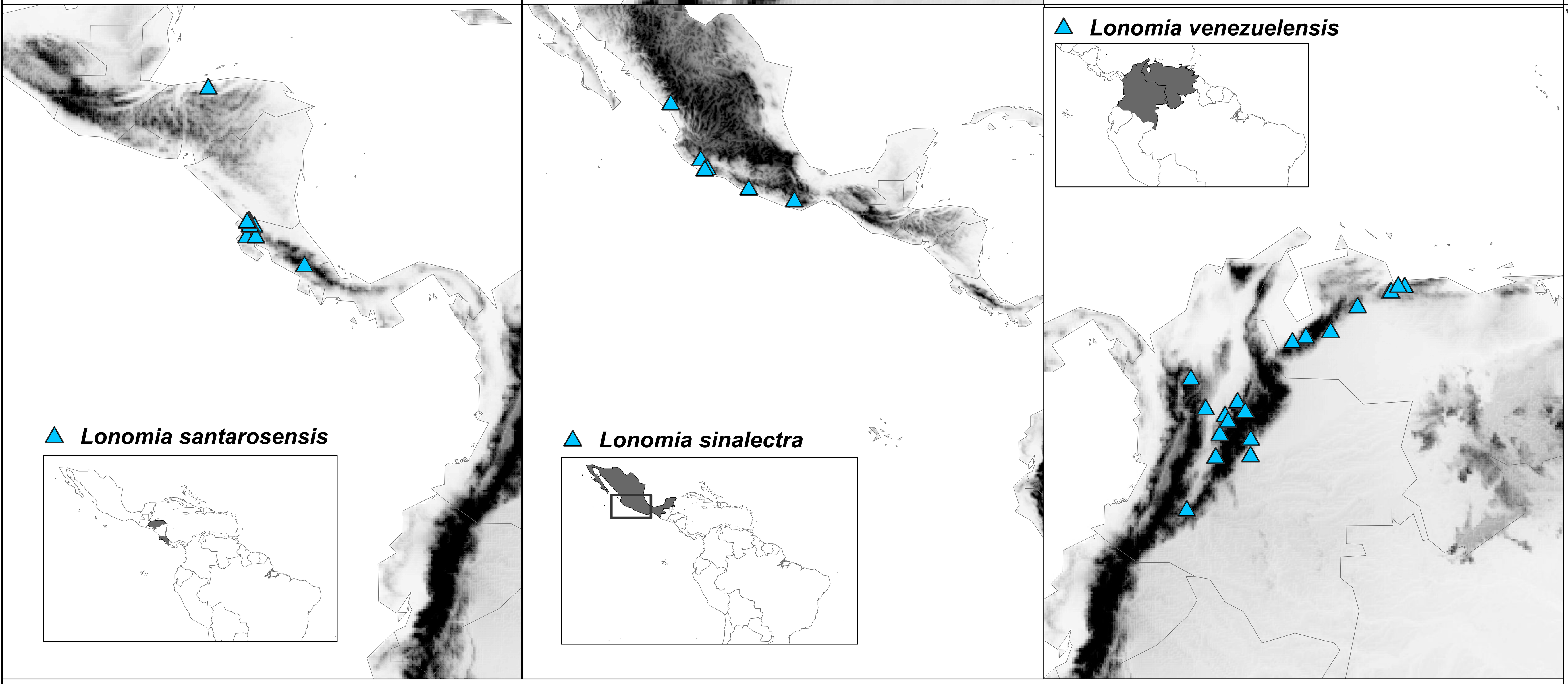


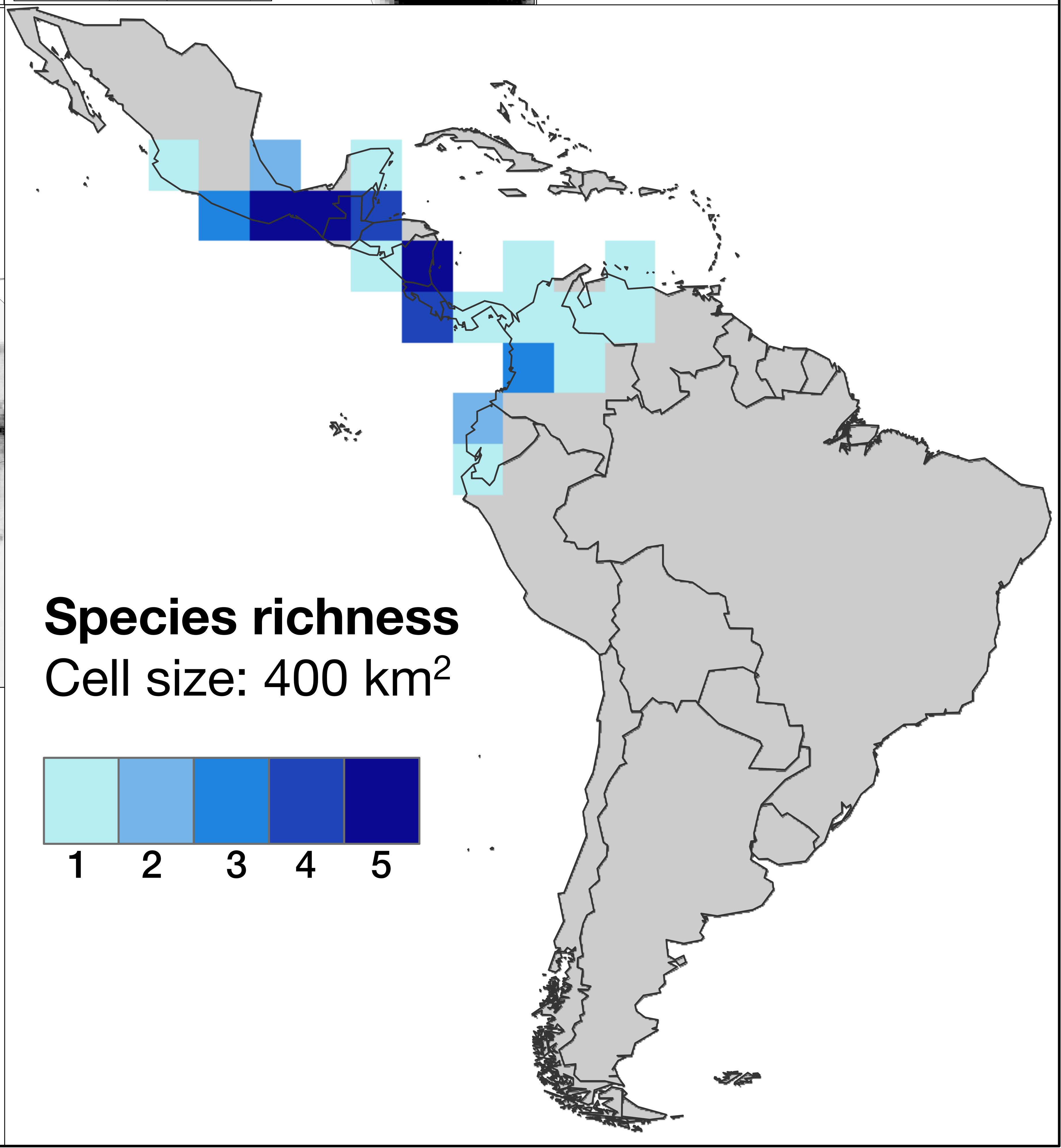


**Figure S5. Electra Group**

This panel shows the distribution of 13 species of *Lonomia* belonging to the electra-group as defined in the present study. The inset maps show the countries where occurrence records are known. The bottom right map shows species richness of the group at a resolution of 400km$^2$ grid cell.
