## Supplementary Table S1 for "Deadly and venomous *Lonomia* caterpillars are more than the two usual suspects"

| Species Group | Species | BIN (as of JUL.15.2022) | Tree# | Authorship | N | N_bc | Typ_bc | Ncountries | Countries | Dmax | DminNN | NN | Notes |
|---|---|---|---|---|---|---|---|---|---|---|---|---|---|
| achelous | *Lonomia achelous* | ADG0261 | **57** | (Cramer, 1777) | 15 | 12 | | 5 | Br, C, FG, G, Su | 0.8 | 1.2 | *L. madrediosiana* | |
| | *Lonomia annamiekae* | AAY4841 | **26** | Brechlin & Meister, 2013 | 2 | 2 | HT | 1 | Pe | 1.8 | 5.2 | *L. viksinyaevi* | |
| | *Lonomia belizonensis* | ACF1945 | **36** | Brechlin *et al.*, 2011 | 25 | 6 | | 1 | Br | 2 | 4.3 | *L. canescens* | |
| | | AAB0791 | | | | 17 | HT | | FG | | | | |
| | *Lonomia beneluzi* | AAK6621 | **22** | Lemaire, 2002 | 10 | 5 | - | 1 | FG | 0 | 6.7 | *L. venezuelensis* | |
| | *Lonomia camox* | AAE0724 | **24** | Lemaire, 1972 | 37 | 27 | HT | 2 | FG, V | 4.2 | 3.3 | *L. canescens* | |
| | *Lonomia canescens* | AAA8506 | **25** | Brechlin & Meister, 2011 | 16 | 12 | HT | 3 | B, E, Pe | 0.8 | 3.3 | *L. camox* | |
| | *Lonomia casanarensis* | AAB4839 | **60** | Brechlin, 2017 | 27 | 25 | HT | 1 | C | 0.3 | 2.7 | *L. diabolus* | |
| | *Lonomia cayennensis* | AAB0796 | **42** | Brechlin & Meister, 2019 | 10 | 10 | HT | 2 | C, FG | 0.6 | 3.7 | *L. CGR01* | |
| | *Lonomia* CGCM01 | AAB4841 | **49** | - | 7 | 7 | - | 1 | Br | 1.2 | 3.7 | *L. sinjaevorum* | |
| | *Lonomia* CGCM02 | ABA7785 | **43** | - | 2 | 2 | - | 1 | Br | 0 | 3.4 | *L. orientoandensis* | |
| | *Lonomia* CGR01 | ACY4181 | **44** | - | 1 | 1 | - | 1 | C | - | 3.7 | *L. cayennensis* | Caterpillar |
| | *Lonomia* CGR02 | ADJ2216 | **48** | - | 1 | 1 | - | 1 | Br | - | 5.2 | *L. sinjaevorum* | Caterpillar |
| | *Lonomia descimoni* | ACR7071 | **37** | Lemaire, 1972 | 39 | 16 | - | 3 | C, E | 2.3 | 2.1 | *L. rubrescens* | |
| | | AAB0795 | | | | 9 | - | | E, Pe | | | | |
| | *Lonomia diabolus* | AAB4840 | **59** | Draudt, 1929 | 27 | 23 | - | 3 | FG, V, TT | 1 | 2.7 | *L. casanarensis* | stat. nov. |
| | *Lonomia francescae* | AAF5434 | **51** | L. Racheli, 2005 | 7 | 2 | - | 1 | E | 3.9 | 2.3 | *L. sinjaevorum* | |
| | | AAF5435 | | | | 4 | - | | | | | | |
| | *Lonomia frankae* | AAF5492 | **27** | Meister *et al.*, 2005 | 16 | 3 | PT | 1 | Pe | 0.2 | 3.9 | *L. pseudobliqua* | |
| | *Lonomia madrediosiana* | AAB4835 | **58** | Brechlin & Meister, 2011 | 36 | 19 | HT | 2 | E, Pe | 1.5 | 1.2 | *L. achelous* | |
| | *Lonomia manabiana* | AED0101 | **32** | Brechlin *et al*., 2013 | 9 | 2 | HT | 1 | E | 0.6 | 1.8 | *L. nigra* | includes *L. oroiana*, syn. nov. (HT with DNA barcode) |
| | *Lonomia maranhensis* | AAB4045 | **56** | Brechlin *et al.*, 2011 | 10 | 10 | HT | 1 | Br | 0.8 | 3.5 | *L. madrediosiana* | |
| | *Lonomia moniqueae* | AAP0951 | **41** | Brechlin & Meister, 2019 | 9 | 6 | HT | 1 | V | 0.4 | 4.8 | *L. diabolus* | |
| | *Lonomia nigra* | AEC7052 | **33** | Brechlin *et al*., 2013 | 8 | 3 | HT | 1 | E | 0.3 | 1.8 | *L. manabiana* | |
| | *Lonomia orientoandensis* | AAB4836 | **45** | Brechlin & Meister, 2011 | 20 | 20 | HT | 3 | C, E, Pe | 2.6 | 3.4 | *L. CGCM02* | |
| | *Lonomia orientocordillera* | AAB4838 | **50** | Brechlin *et al*., 2013 | 22 | 17 | HT | 3 | C, E, Pe | 2.7 | 3 | *L. sinjaevorum* | |
| | *Lonomia panguana* | ACL6324 | **53** | Brechlin, 2017 | 3 | 3 | HT | 1 | Pe | 2 | 2.4 | *L. riojensis* | |
| | *Lonomia parubrescens* | ABZ8294 | **39** | Brechlin & Meister, 2011 | 22 | 3 | | 2 | B, Pe | 2.7 | 1.3 | *L. rubrescens* | |
| | | ACE7193 | | | | 9 | HT | 1 | Pe | | | | |
| | | ACF3136 | | | | 3 | | 1 | Pe | | | | |
| | | ACK9075 | | | | 2 | | 1 | Pe | | | | |
| | *Lonomia pastazana* | AAF5930 | **47** | Brechlin, 2017 | 4 | 4 | HT | 1 | E | 0.6 | 0.5 | *L. siriae* | |
| | *Lonomia pseudobliqua* | AAB0793 | **28** | Lemaire, [1973] | 7 | 4 | - | 1 | Pe | 0.2 | 1.6 | *L. viksinyaevi* | includes *L. vikcuscensis*, syn. nov. (HT with DNA barcode) |
| | *Lonomia puntarenasiana* | AAA7086 | **34** | Brechlin & Meister, 2011 | 35 | 31 | HT | 2 | C, CR, N, P | 2.6 | 1.8 | *L. manabiana* | |
| | *Lonomia rengifoi* | ACS2525 | **23** | Brechlin & Käch, 2017 | 4 | 3 | HT | 2 | C, E | 2.4 | 4.4 | *L. canescens* | |
| | | ACS2526 | | | | 1 | | 1 | E | | | | |
| | *Lonomia riojensis* | AAY4842 | **52** | Brechlin & Meister, 2013 | 2 | 1 | HT | 1 | Pe | - | 2.3 | *L. silkae* | |
| | *Lonomia rubrescens* | ACE5375 | **40** | Brechlin & Meister, 2011 | 25 | 20 | PT | 1 | Pe | 0.5 | 1.3 | *L. parubrescens* | |
| | *Lonomia rubroguyana* | AAB0790 | **38** | Brechlin & Meister, 2019 | 32 | 26 | HT | 1 | FG | 1.8 | 1.8 | *L. parubrescens* | |
| | | ACV3238 | | | | 6 | | 1 | FG | | | | |
| | *Lonomia rufescens* | AAA7087 | **31** | Lemaire, 1972 | 22 | 9 | PT | 2 | C, E | 0.8 | 4.8 | *L. manabiana* | |
| | *Lonomia serranoi* | AAJ2304 | **30** | Lemaire, 2002 | 22 | 13 | HT | 5 | Be, Gt, H, M, S | 2.8 | 5.4 | *L. maranhensis* | includes *L. yucatensis*, syn. nov. (HT with DNA barcode) |
| | *Lonomia silkae* | ABZ8347 | **54** | Brechlin & Meister, 2013 | 5 | 3 | HT | 1 | Pe | 0.5 | 1.7 | *L. sinjaevorum* | |
| | *Lonomia sinjaevorum* | AAB0794 | **55** | Brechlin & Meister, 2011 | 26 | 16 | HT | 1 | B | 3 | 1.7 | *L. silkae* | |
| | | ACF0924 | | | | 6 | PT | 1 | Pe | | | | |
| | *Lonomia siriae* | AAF5930 | **46** | Brechlin & Meister, 2013 | 22 | 14 | HT | 2 | B, Pe | 1.4 | 0.5 | *L. pastazana* | |
| | *Lonomia vanschaycki* | AAB0792 | **35** | Brechlin *et al*., 2013 | 37 | 17 | HT | 3 | C, E, Pe | 0.6 | 4.6 | *L. canescens* | |
| | *Lonomia viksinyaevi* | AAB0793 | **29** | Brechlin & Meister, 2011 | 5 | 5 | HT | 1 | B | 2.2 | 1.6 | *L. pseudobliqua* | |
| electra | *Lonomia columbiana* | AAC9591 | **6** | Lemaire, 1972 | 92 | 11 | HT | 2 | C, E | 3.9 | 2.4 | *L. laalbania* | |
| | | ABY3226 | | | | 10 | | 1 | CR | | | | |
| | | ACG1953 | | | | 4 | | 1 | E | | | | |
| | | ADG2380 | | | | 22 | | 2 | C, P | | | | |
| | *Lonomia concordia* | AAA5428 | **7** | Druce, 1886 | 98 | 59 | LT | 2 | CR, P | 2.6 | 4.3 | *L. quintanarooensis* | includes ssp. *conricana* (HT with DNA barcode) |
| | *Lonomia electra* | ABY6452 | **14** | Druce, 1886 | 47 | 3 | | 1 | M | 2.8 | 1.5 | *L. luteomexicana* | |
| | | ABY6721 | | | | 10 | LT | 1 | Gt | | | | |
| | *Lonomia electra* DHJ02 | AAF5436 | **8** | - | 9 | 9 | - | 1 | CR | 0.3 | 2.9 | *L. quintanarooensis* | |
| | *Lonomia laalbania* | ACO9490 | **4** | Brechlin, 2017 | 2 | 2 | HT | 1 | C | 0 | 2.4 | *L. columbiana* | |
| | *Lonomia luteomexicana* | ABY4571 | **13** | Brechlin & Meister, 2011 | 7 | 2 | | 1 | M | 2 | 1.5 | *L. paralectra* | |
| | | ABY7782 | | | | 3 | | | | | | | |
| | | ACE5178 | | | | 2 | HT | | | | | | |
| | *Lonomia mexilectra* | AAA6679 | **10** | Brechlin & Meister, 2011 | 4 | 4 | HT | 1 | M | 0.8 | 2.6 | *L. sinalectra* | |
| | | ABY8855 | | | 5 | 4 | | | | 1.8 | | | This BIN recognized as ssp. m*expueblensis* (HT with DNA barcode) |
| | *Lonomia minca* | ACT2303 | **1** | Brechlin, 2017 | 7 | 7 | HT | 1 | C | 0.2 | 4.8 | *L. venezuelensis* | |
| | *Lonomia paralectra* | AAA6675 | **12** | Brechlin & Meister, 2011 | 17 | 16 | HT | 1 | Gt | 2.3 | 1.5 | *L. luteomexicana* | |
| | | ADL2612 | | | | 1 | | 1 | M | | | | |
| | *Lonomia peggyae* | AAC9590 | **5** | Brechlin *et al*., 2013 | 39 | 6 | | 1 | E | 2.9 | 2.4 | *L. columbiana* | |
| | | ACE5050 | | | | 14 | HT | | | | | | |
| | | ACF7094 | | | | 1 | | | | | | | |
| | *Lonomia quintanarooensis* | AAA6676 | **9** | Brechlin & Meister, 2011 | 33 | 23 | HT | 5 | Be, Gt, H, M, N | 2.1 | 2.9 | *L. electraDHJ02* | includes ssp. q*uinchiapasiana* (HT with DNA barcode) |
| | *Lonomia santarosensis* | AAA7085 | **3** | Brechlin & Meister, 2013 | 27 | 23 | HT | 3 | CR, H, N | 2.2 | 3.5 | *L. columbiana* | |
| | *Lonomia sinalectra* | AAA6678 | **11** | Brechlin & van Schayck, 2015 | 3 | 16 | HT | 1 | M | 4.3 | 2.6 | *L. mexilectra* | |
| | | AAA6680 | | | 1 | | | | | | | | |
| | | ABA7784 | | | 3 | | | | | | | | |
| | | ABY6181 | | | 2 | | | | | | | | |
| | | ABY6182 | | | 2 | | | | | | | | |
| | | ACE3733 | | | 2 | | | | | | | | |
| | | ACF3079 | | | 2 | | | | | | | | |
| | | ACZ2305 | | | 1 | | | | | | | | |
| | *Lonomia venezuelensis* | AAC1812 | **2** | Lemaire, 1972 | 34 | 29 | - | 2 | C, V | 0.9 | 3.5 | *L. laalbania* | |
| obliqua | *Lonomia antoniae* | ABA3772 | **15** | Brechlin & Meister, 2015 | 31 | 11 | HT | 1 | Br | 1.4 | 5.2 | *L. obliqua* | |
| | *Lonomia* CGCM03 | ABA7783 | **16** | - | 3 | 3 | - | 1 | Br | 0.2 | 1.3 | *L. leopoldina* | |
| | *Lonomia leopoldina* | AAC1813 | **17** | Brechlin & Meister, 2011 | 9 | 4 | HT | 1 | Br | 2.6 | 1.3 | *L. CGCM03* | |
| | | ACV8217 | | | | 1 | | | | | | | |
| | *Lonomia obliqua* | AAB4042 | **20** | Walker, 1855 | 49 | 33 | - | 3 | B, Br, Pa | 1.8 | 1.8 | *L. parobliqua* | |
| | *Lonomia parobliqua* | AAB4043 | **21** | Brechlin *et al.*, 2011 | 37 | 22 | HT | 3 | A, Br, GF | 2.2 | 1.8 | *L. obliqua* | |
| | | ACE6785 | | | | 12 | | 4 | A, B, Br, Pa | | | | |
| | *Lonomia* RRBR01 | AAB4044 | **18** | - | 13 | 9 | - | 1 | Br | 1 | 6.7 | *L. obliqua* | |
| | *Lonomia rufobahiana* | ABA3773 | **19** | Brechlin & Meister, 2013 | 4 | 3 | HT | 1 | Br | 0.3 | 3.9 | *L. obliqua* | |
